# MeC3HDZ1/MeCNA is a strong candidate for cassava storage root productivity improvement

**DOI:** 10.1101/2023.03.07.531519

**Authors:** Anna Solé-Gil, Anselmo López, Damiano Ombrosi, Cristina Urbez, Javier Brumós, Javier Agustí

## Abstract

The storage root (SR) of cassava is the main staple food in sub-Saharan Africa, where it feeds over 500 million people. However, little is known about the genetic and molecular regulation underlying its development. Unraveling such regulation would pave the way for biotechnology approaches aimed at enhancing cassava productivity. Anatomical studies indicate that SR development relies on the massive accumulation of xylem parenchyma, a cell-type derived from the vascular cambium. The C3HDZ family of transcription factors regulate cambial cells proliferation and xylem differentiation in Arabidopsis and other species. We thus aimed at identifying C3HDZ proteins in cassava and determining whether any of them shows preferential activity in the SR cambium and/or xylem. Using phylogeny and synteny studies, we identified eight C3HDZ proteins in cassava, namely MeCH3DZ1-8. We observed that the expression of *MeC3HDZ1* in SR cambium and xylem is higher than that of any other *MeC3HDZ* gene in any of the SR vascular tissues or any of the other vegetative organs. We established an *in-silico* pipeline which revealed the existence of a number of theoretical C3HDZ targets displaying significant preferential expression in the SR. Subsequent Y1H analyses proved that MeC3HDZ1 can bind canonical C3HDZ binding sites in the promoters of these targets. Transactivation assays demonstrated that MeC3HDZ1 can regulate the expression of genes downstream of promoters harboring such binding sites, thereby demonstrating that MeC3HDZ1 is a C3HDZ transcription factor which constitutes a strong candidate for future biotechnology strategies directed at increasing cassava productivity.

## INTRODUCTION

The current world’s population has increased three-fold in the last 70 years (www.un.org), reaching over eight billion people. Predicting models indicate that, by the end of the 21^st^ century, it will reach ten billion. Developing countries are experiencing the largest population increase and this growth patterns are predicted to continue over the next decades. Considering the climate change context and the decrease in arable land that population growth entails, quick and targeted improvement of our main staple crops production to ensure food security is a top priority.

Cassava (*Manihot esculenta*) is the fifth staple crop worldwide and the top one in sub-Saharan Africa, where it currently feeds over five hundred million people (www.fao.org). Due to a variety of reasons, cassava is the preferred crop for self-sustained farmers in many sub-Saharan Africa regions. First, cassava displays strong resistance to high temperatures and drought. Second, it presents an outstanding capacity to accumulate carbohydrates in the form of starch in its storage root (from now on SR), the edible part of the plant. Third, the SR remains in good conditions in the soil for a long time after reaching its full development, allowing for gradual harvest depending on needs. Fourth, cassava can be clonally propagated in an extremely easy manner (Lebot V, 2009).

In the last couple of decades, strong research efforts have been made to improve cassava in terms of SR nutritional quality, disease resistance and postharvest deterioration (Sayre et al., 2011; Otun et al., 2022; Shakir et al., 2022). Remarkably, little is known about cassava root development. Indeed, although reports on the anatomical dynamics on SR formation exist(Chaweewan and Taylor, 2015), the molecular mechanisms underlying SR development are poorly understood (Vanderschuren and Agusti, 2022).

The mature cassava root system consists of two types of roots that coexist, namely the fibrous root (FR) and the SR (Lebot V, 2009). During the first steps of root development, all roots are FR. Later on, through signals that are currently unknown, a subset of these roots experiences several developmental changes to become SRs (Alves AAC, 2002; El-Sharkawy, 2004). Histologically, the differentiation of the two types of roots can be explained by the type of xylem that they develop. Whereas FRs develop xylem vessels and fibers, SRs develop, almost exclusively, xylem parenchyma (Vanderschuren and Agusti, 2022). Xylem vessels are the major water transporters, thus supporting a strong water pressure (Esau, 1961). The fibers’ role is mainly complementary to that of vessels, providing mechanical support (Dickison, 2000). For these reasons, vessels and fibers develop thick secondary cell walls that are strongly lignified (Ruzicka et al., 2015). The xylem parenchyma cells do not develop lignified secondary cell walls (Esau, 1961). Instead, these xylem cells possess the ability of synthesizing and storing starch (Esau, 1961; Ruzicka et al., 2015). Considering that xylem cells develop from the cambial meristematic cells, undergoing several maturing steps that lead them to acquire their final identity (Agustí and Blázquez, 2020), understanding how cassava cambial cells proliferate and how developing xylem cells acquire the parenchymatic identity, can tremendously help improving cassava productivity.

Forward genetic approaches aiming at identifying root developmental regulators in cassava are scarce. Partially, this is due to the cassava adult plant’s size, the relatively long flowering time, and the fact that the storage root takes several weeks to develop. However, the genetics of root development -including that concerning cambium activity and xylem differentiation-are well documented in the model species *Arabidopsis thaliana*, making it possible to perform gene-targeted approaches aimed at understanding the molecular regulation of root development in cassava based on such knowledge. Notably, our previous studies succeeded in identifying orthologues in cassava of Arabidopsis regulators of vessels or fibers differentiation displaying differential expression patterns between the FR and the SR (Siebers et al., 2017). To take a step forward, here we aimed at identifying genetic regulators of the vascular pattern establishment and xylem identity acquirement in cassava. In Arabidopsis, the C3HDZ family of transcription factors has been shown to play a key role in this respect(Ohashi-Ito and Fukuda, 2003; Ohashi-Ito et al., 2005; Prigge and Clark, 2006; Carlsbecker et al., 2010; Smetana et al., 2019). The C3HDZ family consists of five redundantly acting paralogues in Arabidopsis, namely: PHABULOSA (PHB), REVOLUTA (REV), PHAVOLUTA (PHV), CORONA (CNA) and ARABIDOPSIS THALIANA HOMEOBOX 8 (AtHB8) (Baima et al., 2001; Carlsbecker et al., 2010). Consistently with the reported redundancy, most of the gene s encoding these proteins show overlapping expression patterns (Carlsbecker et al., 2010). In general, the *AtC3HDZ* genes are expressed in the cambial zone. Among them, *AtHB8* expresses preferentially in the xylem side of the cambium, supporting the idea that C3HDZ proteins mark cells with xylem identity (Baima et al., 2001; Carlsbecker et al., 2010; Smetana et al., 2019). Genetic and molecular analyses have helped fine-tuning the function of the C3HDZ proteins, demonstrating that, although a general redundancy exists, each member of the family may play dominant roles in specific developmental processes. For example, *rev* loss of function mutants display a lack of interfascicular fibers in the stem (Zhong and Ye, 1999; Prigge and Clark, 2006). Also, plants ectopically expressing *AtHB8* show enhanced secondary growth (Baima et al., 2001), and the *CORONA* gain of function *icu-4* mutant develops more vascular tissue than WT (Ochando et al., 2008). However, single *Atc3hdz* loss of function mutants show no obvious xylem or cambium phenotype; defects in this respect being only found in high order mutants (Carlsbecker et al., 2010). Thus, quadruple mutants show some xylem defects and scattered cambial divisions, and mutants for all five genes display no xylem development (Carlsbecker et al., 2010; Smetana et al., 2019). All in all, these data reinforce the idea that the C3HDZ family is required for cambium activity and xylem establishment and that strong redundancy exists between the members of the family in the regulation of such developmental processes in Arabidopsis. The *C3HDZ* expression levels are negatively regulated by the *miRNA165/166* (Rhoades et al., 2002; Reinhart et al., 2002; Emery et al., 2003; Tang et al., 2003). The differential presence of *miRNA165/166* imposes high or low accumulation of *C3HDZ* transcripts (Emery et al., 2003; Carlsbecker et al., 2010). High *miRNA165/166* accumulation leads to protoxylem differentiation, while low accumulation results in metaxylem differentiation, demonstrating a fundamental role for the family in the determination of xylem cell-type identity (Carlsbecker et al., 2010).

Characterization of C3HDZ TFs in *Zinnia elegans* and rice demonstrated that the essential functions of the family in vascular development are most likely conserved across species (Ohashi-Ito and Fukuda, 2003; Ohashi-Ito et al., 2005; Itoh et al., 2008). Importantly though, in *Populus*, the C3HDZ effect on vascular cambium morphogenesis seems to be mainly capitalized by one single protein, namely PopREV (Robischon et al., 2011), the Populus orthologue of REVOLUTA. This result implies that the observed redundancy in *Arabidopsis* may not equally apply to all species. Thus, in this work we aimed at identifying C3HDZ proteins preferentially operating in the storage root and potentially regulating developmental aspects of SR formation such as cambium activity and xylem parenchyma identity acquisition. To that end, we first used phylogeny and synteny to determine the cassava *Manihot esculenta*) orthologues of the C3HDZ proteins. Using gene expression analyses, we determined that *MeC3HDZ1*, a member of the *MeC3HDZ* family encoding a CORONA orthologue, not only displays preferential expression in the storage root in comparison with the rest of vegetative organs but also shows a much stronger expression in the storage root cambium and developing xylem than any of the other members of the family. Furthermore, we have demonstrated that MeC3HDZ1 can bind to promoters carrying the canonical C3HDZ binding site and regulate the expression of genes downstream of such promoters, indicating that MeC3HDZ1 is able to carry out C3HDZ transcription factor activity. We propose that MeC3HDZ1 may be relevant both for the regulation of xylem parenchyma identity acquisition and for the proliferation of cambium cells during cassava SR development, implying that it can be exploited for future biotechnology approaches aimed at improving cassava productivity.

## RESULTS

### Cassava possesses eight C3HDZ orthologues in its genome

To identify the most likely orthologues in cassava for each of the five C3HDZ transcription factors that exist in Arabidopsis, we first constructed maximum likelihood phylogenetic trees using potential C3HDZ orthologues identified in eleven more species (see Methods and Materials for details). Our BLAST search identified 89 amino acid sequences throughout the twelve species (Supplemental File 1). The tree displayed five different clades, among which we identified eight potential cassava C3HDZ proteins (Figure 1), from now on MeC3HDZ1 to 8. Clades were named depending on the Arabidopsis C3HDZ protein/s that they contained. While MeC3HDZ1 and 2 clustered in the CORONA (CNA) clade, MeC3HDZ3 and 4 fell in the AtHB8 clade and the REVOLUTA (REV) clade contained MeHDZIP5 and 6. MeC3HDZ7 and 8 were found in the PHABULOSA/PHAVOLUTA (PHB/PHV) clade. Identifying two cassava orthologues per Arabidopsis protein is a quite common feature, since cassava experienced a genomic duplication over evolution (Prochnik et al., 2012). The fifth clade was termed ATHB8/CNA because it located between such clades (Figure 1). It only contained monocot (rice or foxtail millet) sequences. Synteny analyzes confirmed that cassava genes encoding proteins that fall within the CNA, AtHB8 or REV clades are orthologues of such genes and that MeC3HDZ7 and MeC3HDZ8 could be orthologues either of PHB or PHV (Supplemental figure 1).

**Figure 1.**
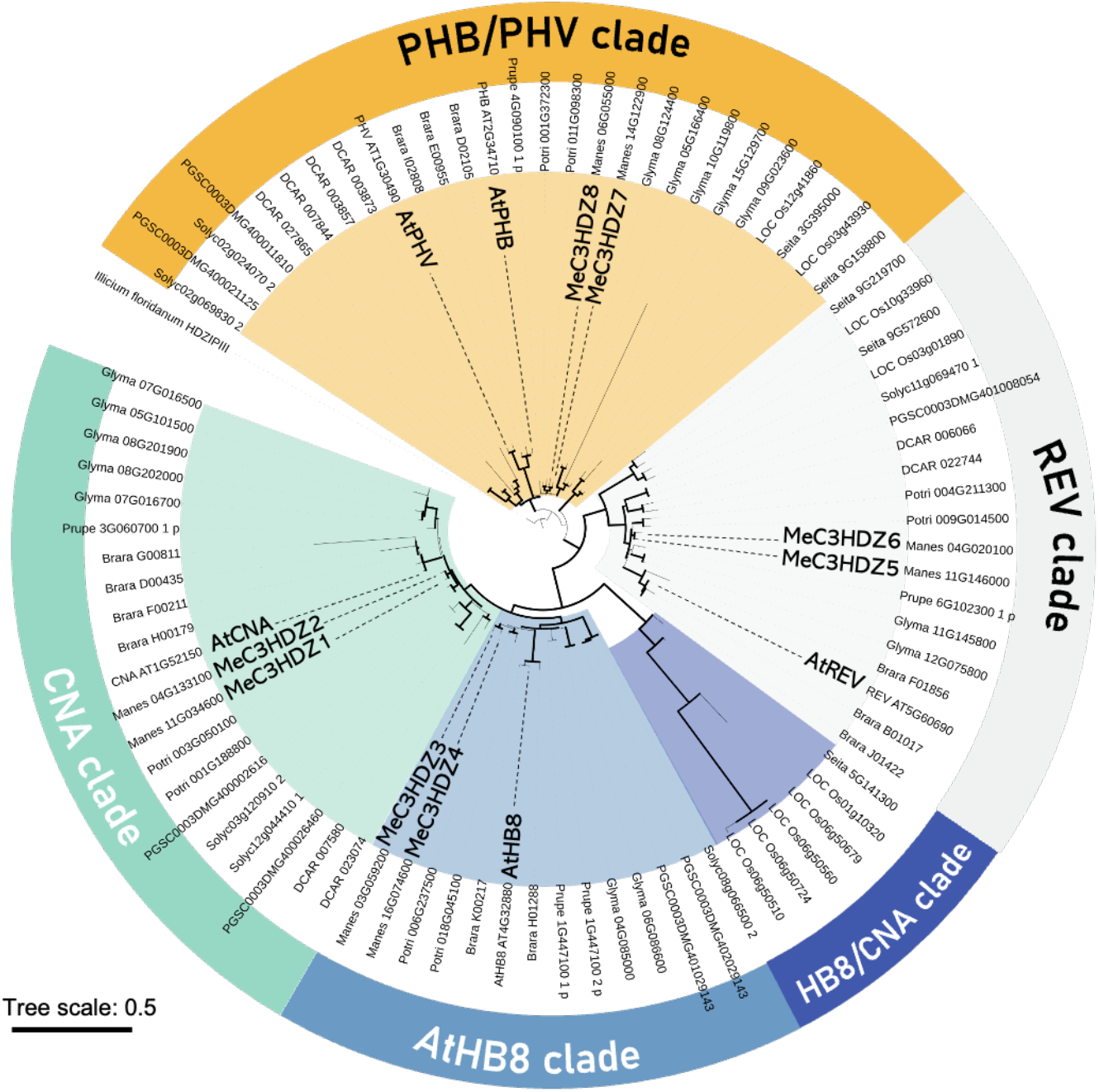
Identification of putative C3HDZ proteins in cassava. Figure 1. Identification of putative C3HDZ proteins in cassava. Maximum likelihood tree generated with protein sequences of: cassava (*Manihot esculenta*; proteins code: Manes), Arabidopsis (*Arabidopsis thaliana*; proteins code: AT), field mustard (*Brassica rapa*; proteins code: Brara), tomato (*Solarium Lycopersicon*; proteins code: Solyc), potato (*Solarium tuberosum*; proteins code: PGSC), carrot(*Daucus carota*; DCAR), soybean (*Glycine max*; proteins code: Glyma), black cottonwood (*Populus trichocarpa*; proteins code: Potri), peach (*Prunus persica*; proteins code: Prupe), rice (*Oryza sativa)*, foxtail millet (*Setaria italica*; proteins code: Seita) and purple anise (*llicium floridanum*; ANA clade, outgroup). Clades are defined based on Arabidopsis proteins as follows: yellow PHB/PHV, grey REV, dark blue HB8/CNA monocot (rice and setaria), light blue AtHB8, turquoise CNA. Arabidopsis and cassava orthologues are marked. Branch support values are based on Shimodaira-Hasegawa-like approximate likelihood ratio test (SH-like aLRT), and values over 0.8 are marked with thicker black branches.

### *MeC3HDZ1* displays preferential expression in the storage root cambium and developing xylem

Considering the relevant role of the C3HDZ proteins in the establishment of the vascular patterns, cambium activity and xylem development (Baima et al., 2001; Carlsbecker et al., 2010), we searched for *MeC3HDZ* genes preferentially expressed in SR in comparison to FR, as those would be good candidates for targeted biotechnology manipulation of the SR development. Our expression analyses revealed that all *MeC3HDZ* genes except *MeC3HDZ6* displayed enhanced expression in SR in comparison to FR (Figure 2A). *MeC3HDZ8*, *MeC3HDZ5* and *MEC3HDZ1* displayed the highest differential expression in comparison to FR, with 3.737-fold, 3.167-fold and 2.933-fold, respectively (Figure 2A, Supplemental table 1). To understand the expression profile of the *MeC3HDZ* genes in the rest of vegetative (aerial) organs, we checked their expression in adult leaves, stems and sprouts. The expression of each gene was referred to that found in FR to generate a comparative context with our SR differential expression analyzes. In adult leaves all genes displayed considerably less expression levels than in FR (Figure 2A, Supplemental table 1). In stems, all genes displayed a similar expression level to that found in FR, except for *MeC3HDZ5*, which showed 3,019-fold expression (Figure 2A, Supplemental table 1). In the case of sprout, *MeC3HDZ1*, *MeC3HDZ2* and *MeC3HDZ6* showed weak expression levels, whereas the rest of the genes expressed at similar levels as in FR (Figure 2A, Supplemental table 1). In conclusion, our experiments revealed that, in general, all *MeC3HDZ* genes express at higher levels in SR than in any other vegetative organ of the plant. Our data indicated that *MeC3HDZ8*, *MeC3HDZ5* and, to a lesser extent, *MeC3HDZ1*, display the highest differential expression between SR and FR (Figure 2A), suggesting that these genes may be relevant for SR targeted biotechnology applications. However, such differential expression does not only depend on the expression levels in the SR but also on those in the FR. In this regard, the expression levels of *MeC3HDZ5* and *MeC3HDZ8* are very much reduced in FR in comparison with the expression levels of *MeC3HDZ1*, the gene displaying highest expression levels in this organ (Figure 2A).

**Figure 2.**
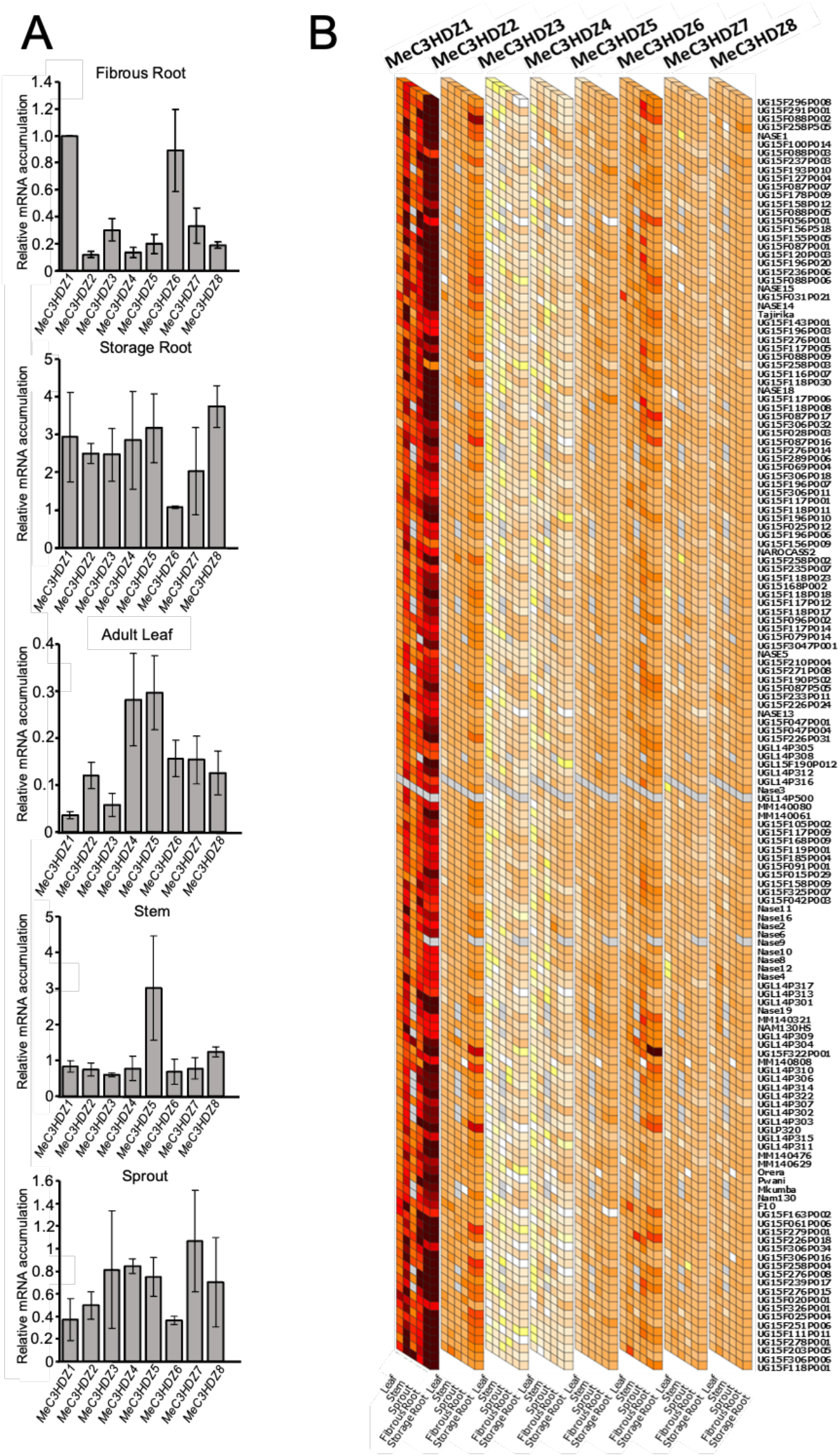
Relative mRNA accumulation of the *MeC3HDZ* genes in different plant organs. (A) Quantitative RT-PCR analyses revealed the relative mRNA accumulation of each *MeC3HDZ* gene in fibrous root, storage root, adult leaf, stem or sprout. For fibrous root, mRNA accumulation levels were normalized to that of *MeC3HDZ1*. For the rest of organs, mRNA accumulation of each gene was normalized to that of fibrous root. (B) Expression levels of the eight *MeC3HDZ* genes in different plant organs across 150 cassava accessions. Diagram generated from data contained in Ogbonna et al. (2021).

To determine what *MeC3HDZ* gene could be our best candidate for biotechnology applications, we reasoned that it is worth considering that there is a large number of cassava accessions/cultivars and that different accessions may behave differently in terms of differential gene expression between different organs. Therefore, we made use of a RNAseq-based transcriptomic atlas for 150 cassava accessions (Ogbonna et al., 2021) to check the expression levels of the eight *MeC3HDZ* genes in all vegetative organs in all such accessions. Our results revealed that in almost all accessions *MeC3HDZ1* is, by far, the gene displaying the highest level of expression in the SR (Figure 2B), and that it expresses in higher levels in the SR than in the rest of the organs.

Taken all together, our data argue for *MeC3HDZ1* as a highly valuable candidate for future biotechnology strategies aimed at maximizing the SR developmental capacities and, therefore, cassava productivity. Nevertheless, since such traits rely on cambium activity and xylem parenchyma proliferation, we decided to check the *MeC3HDZ* gene family expression levels in such tissues. To maintain a comparative context with our previous analyzes, we compared the expression of the eight genes in cambium or xylem with that in FR. Our results revealed that, indeed, *MeC3HDZ1* is the gene displaying the highest expression level in cambium, displaying a 4.369-fold expression in comparison to that in FR (Figure 3A; Supplemental table 1). As for the SR xylem, three genes showed enhanced expression in comparison to FR, namely *MeC3HDZ8* (2,604-fold), *MeC3HDZ4* (2,429-fold) and *MeC3HDZ1* (2,356-fold) (Figure 3B; Supplemental table 1). However, considering the low expression of *MeC3HDZ8* and *MeC3HDZ4* in FR (Figure 2A), *MeC3HDZ1* seems to be the highest expressed one in such tissue too. To obtain a complete picture of the *MeC3HDZ* gene family expression in the SR vascular tissues, we expanded our analyzes to phloem, where all genes displayed reduced expression in comparison to that found in FR (Figure 3C; Supplemental table 1). In view of the obtained results, we reinforced our conclusion that *MeC3HDZ1* is a highly valuable gene for SR biotechnology application. Thus, we focused on that gene for further experimentation.

**Figure 3.**
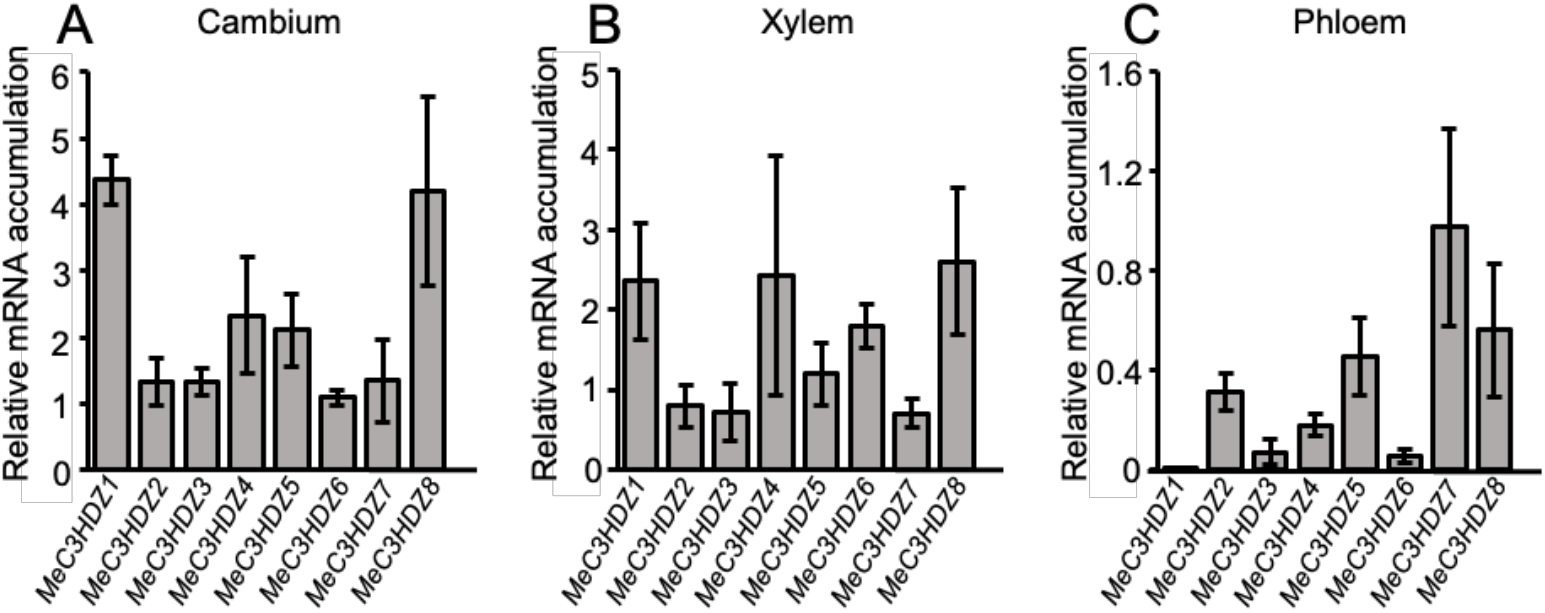
Relative mRNA accumulation of the *MeC3HDZ* genes in storage root vascular tissues. Quantitative RT-PCR analyses revealed the relative mRNA accumulation of each *MeC3HDZ* gene in storage root cambium (A), storage root xylem (B) or storage root phloem (C). The mRNA accumulation of each gene was normalized to that of fibrous root.

### MeC3HDZ-1 possesses C3HDZ activity

After having identified *MeC3HDZ1* as the *MeC3HDZ* with highest expression in SR cambium and xylem, we aimed at determining whether MeC3HDZ1 possesses C3HDZ activity. To that end, we decided to test whether MeC3HDZ1 can both bind the promoter of theoretical C3HDZ targets and regulate their expression. To put our experimentation in the context of SR, we decided to use as theoretical target a gene showing significant preferential expression in SR in comparison to FR. To that end, we first selected 245 genes displaying the strongest significant preferential expression in SR in comparison to FR making use of previously published transcriptomic data (Wilson et al., 2017) (Supplemental table 2; Supplemental Figure 2). We then obtained, for each gene, up to the first 1000 bp of their promoters (i.e the first 1000 bp upstream of the first ATG or the maximum length when the promoter region was less than 1000bp; Supplemental file 2; Supplemental Figure 2). We used JASPAR (jaspar.genereg.net) (Castro-Mondragon et al., 2022) to look for position frequency matrices for *CORONA*, which binding sites happen to be the C3HDZ canonical one (GTAAT(G/C)AT(T/G)(A/G)C; Figure 4A; Supplemental Figure 2). We used the canonical C3HDZ binding sites associated matrices and the 245 promoter sequences to run the web tool MORPHEUS (Minguet et al., 2015), a software providing scores to promoters according on their likeliness to be bound by a given transcription factor depending on the number of repetitions of the transcription factor binding site and the proximity of such sites to the first ATG. With this search, we identified the promoter of *Manes.06G121400* as our best candidate for further experimentation (Figure 4B; Supplemental file 3; Supplemental Figure 2). The workflow for the identification of the *Manes.06G121400* promoter is summarized in Supplemental Figure 2. It is worth mentioning that *Manes.06G121400* expression is clearly enriched in the storage root (Supplemental Figure 3). Remarkably, we used the PlantPAN 3.0 database (Chow et al., 2018)providing genome visualization of ChIP-seq data, to analyze the promoter of the *Manes.06121400* Arabidopsis orthologue *AT5G63530*. Results revealed that, as it is the case for the *Manes.06121400* promoter, the *AT5G63530* promoter harbors several C3HDZ binding sites close to the initiation of transcription site (Supplemental Figure 4).

**Figure 4.**
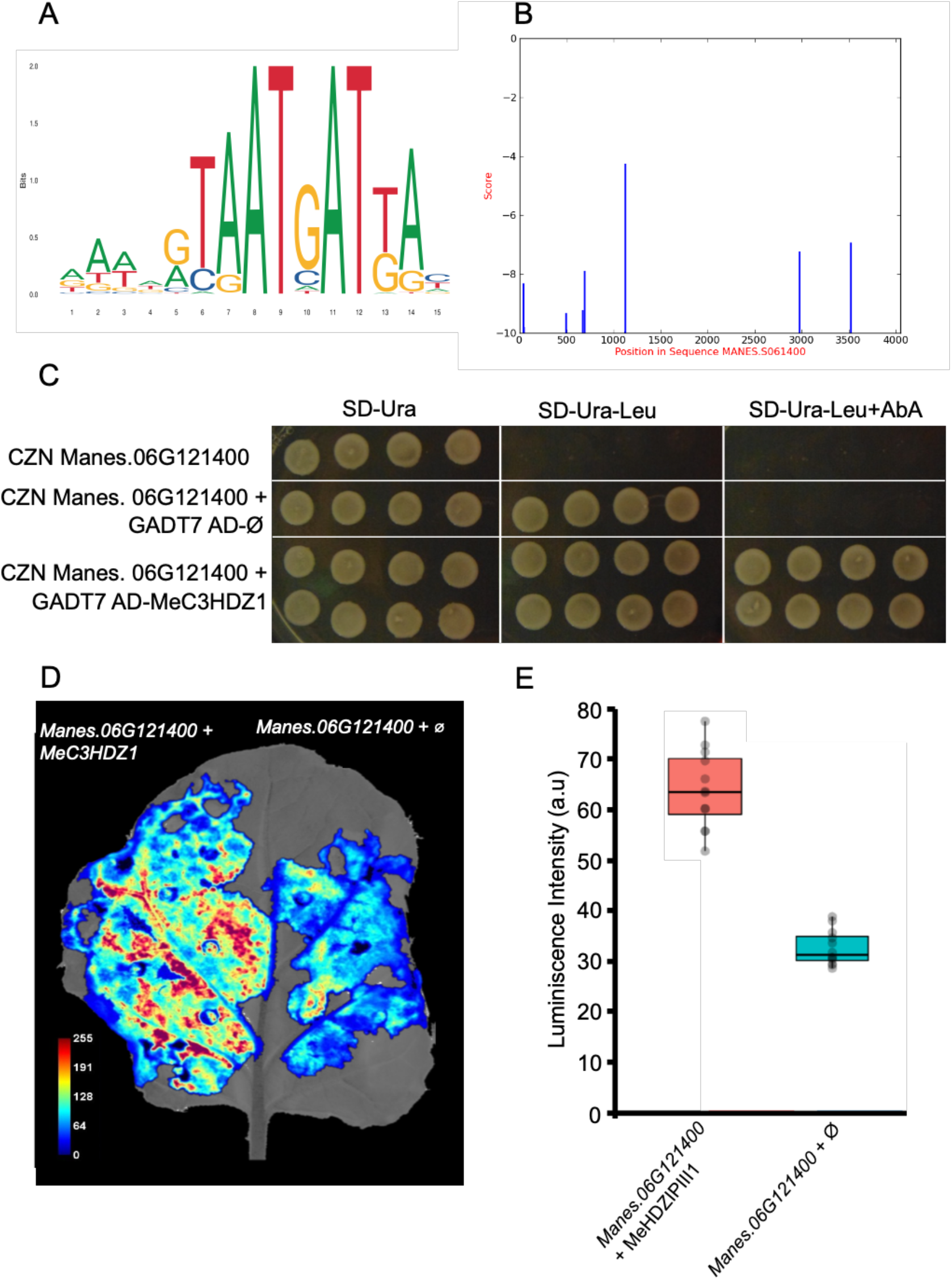
MeC3HDZ1 possesses C3HDZ transcription factor activity. (A) MEME representing the canonical C3HDZ binding site. (B) Canonical C3HDZ binding sites with high probability for C3HDZ binding are present in the *Manes.06G121400* promoter. (C) Y1H analyses show that C3HDZ1 can bind to the *Manes.06G121400* promoter. (D-E) Transactivation assays demonstrate that C3HDZ1 can regulate the transcription downstream of the *Manes. 06G121400* promoter.

Next, we tested the capacity of MeC3HDZ1 to bind the promoter of *Manes.06G121400*. To that end we performed a yeast one-hybrid assay. Since the first 500bp of the *Manes.06G121400* promoter contain two C3HDZ binding sites (Figure 4B; Supplemental file 3), we cloned this sequence in the bait construct. In parallel, the full-length *MeC3HDZ1* coding region was cloned next to the activation domain (AD) in the GADT7 prey vector. We transformed yeast either with the bait construct harboring the *Manes.06G121400* promoter. After confirming that autoactivation of the promoter did not take place, the yeast containing the bait was transformed with either the empty prey vector or with the prey vector expressing *MeC3HDZ1*. Our results confirmed that, indeed, MeC3HDZ1 can bind the *Manes.06G121400* promoter (Figure 4C; Supplemental Figure 5). To test whether MeC3HDZ1 can bind and regulate the transcription of genes downstream of promoters containing C3HDZ binding sites *in planta*, we performed a luciferase transactivation assay in *Nicotiana benthamiana* leaves. Leaves infiltrated only with the *pManes.06G121400::LUC* construct displayed residual *LUC* luminiscence, probably due to the activity of the *N.benthamiana* native C3HDZ proteins (Figure 4D and E). However, when infiltrated together with the *35S::VP16-MeC3HDZ1* we observed more than two-fold *LUC* expression activity increase, confirming that MeC3HDZ1 can bind and regulate the expression of genes containing C3HDZ binding sites in their promoter (Figure 4D and E). Our results, thus, demonstrate that MeC3HDZ1 displays C3HDZ transcription factor activity.

## DISCUSSION

Understanding the molecular mechanisms underlying cassava SRs development is of paramount relevance for downstream biotechnological applications. Identifying genes regulating cambium activity and/or xylem parenchyma determination and proliferation in a preferential manner in SRs represents a first step to implement approaches specifically targeted to enhancing the development of such roots and, thus, cassava productivity. In this respect genes operating in upstream positions in regulatory networks for these biological processes are, probably, the most promising candidates. *MeC3HDZ1* fulfills all these criteria. First, C3HDZ proteins have contrasted activity in cambium activity and xylem development (Baima et al., 2001; Carlsbecker et al., 2010; Smetana et al., 2019). Second, *MeC3HDZ1* encodes a *CORONA* orthologue, and the *CORONA* gain of function mutant *icu4-1* has been shown to stimulate the production of vascular tissues (Ochando et al., 2008). Third, *MeC3HDZ1* is expressed in considerably higher levels in SR cambium and xylem than any of the other *MeC3HDZ* genes in any other vegetative organ (including the FR). Fourth, this preferential expression in the storage root is conserved in, at least, 150 cassava accessions (Figure 2B) and, probably, in many other. Fifth, we have demonstrated that MeC3HDZ1 possesses C3HDZ activity. Further genetic approaches are needed to confirm that, as it is the case for Arabidopsis C3HDZ proteins in root development, MeC3HDZ1 also plays a pivotal role in cambium activity and xylem development in cassava SR. In this respect, it will be interesting to analyze *MeC3HDZ1* CRISPR mutant lines, as well as lines overexpressing the gene. Conditional lines expressing only at will *miRNA165/166*, reportedly targeting *C3HDZ* transcripts (Rhoades et al., 2002; Reinhart et al., 2002; Emery et al., 2003; Tang et al., 2003), *35S::MeC3HDZ1* or a *miRNA165/166*-resistant versions of *MeC3HDZ1* would also help understanding whether manipulating *MeC3HDZ1* expression levels on demand can be of potential use to control SR development in a more efficient and precise manner by aiming at generating specific root sizes.

We have established an *in-silico* pipeline to identify theoretical MeC3HDZ targets. Employing this approach, which essentially could be implemented for any transcription factor provided that the binding sites are known, we have identified *Manes.06G121400*. This gene is expressed in a significant preferential manner in the SR (Supplemental Figure 4), it would be possible to localize specific signals to the SR cambium and xylem by expressing genes under the control of *Manes.06G121400* promoter. For example, expressing a *miRNA165/166* resistant version of *MeC3HDZ1* under the *Manes.06G121400* promoter would provide the opportunity to enhance the C3HDZ function specifically -or in a significantly more prominent manner-in the SR. In the same way, expressing the *miRNA165/166* under the *Manes.06G121400* promoter may help understanding the effect of minimizing the C3HDZ function specifically in the SR.

Understanding what signals regulate *MeC3HDZ1* expression will also be very informative. Hormones, such as auxin, have been shown to regulate the expression of *C3HDZ* genes in Arabidopsis and other species (Baima et al., 1995; Ohashi-Ito and Fukuda, 2003) thus being likely that this is also the case in cassava. Considering that the SR of cassava develops high amounts of xylem parenchyma and synthesizes (and stores) high amounts of starch, it is a good model to understand whether some signals such as sugar are important for the regulation of C3HDZ activity during SR development. It would also be valuable to unravel the effect of soil conditions like pH, drought, salinity or compaction levels on *MeC3HDZ1* expression and or MeC3HDZ1 activity.

To date, very little is known about the molecular mechanisms underlying the cell type specification of xylem parenchyma in plants, perhaps because in Arabidopsis this cell type is difficult to access. However, the relevance of this tissue is clear from an agronomical point of view. Thus, research in cassava SR, where xylem parenchyma is very prominent, may provide new concepts on the fundamental aspects of xylem parenchyma biology. In this way, assessing a function for C3HDZ proteins in xylem parenchyma development would be of great importance not only for applied purposes but also for elementary biology conceptual progress.

In summary, we have identified a protein possessing C3HDZ activity in cassava which displays high and preferential expression in SR cambium and xylem. Due to its nature and reported regulatory activity of its orthologues in Arabidopsis and other species in terms of cambium activity and xylem development, C3HDZ1 holds great promise for future biotechnology approaches. Considering the widespread preferential expression of *MeC3HDZ1* in the SR across a large number of cassava cultivars, any biotechnology application with this gene will be highly relevant, as it could be translated into virtually any cassava cultivar.

## METHODS

### Construction of phylogenetic trees and synteny analyses

To construct the C3HDZ phylogenetic tree (Figure 1; Supplemental File 1) we first search for potential C3HDZ sequences in the cassava (*Manihot esculenta*), black cottonwood (*Populus trichocarpa*), field mustard (*Brassica rapa*), potato (*Solanum tuberosum*), foxtail millet (*Setaria italica*), tomato (*Solanum lycopersicum*), peach (*Prunus persica*), rice (*Oryza sativa*), soybean (*Glycine max*) and carrot (*Daucus carota*) genomes. To that end we used the amino acid sequences of the *Arabidopsis thaliana* C3HDZ proteins - namely REVOLUTA/AT5G60690, PHABULOSA/AT2G34710, PHAVOLUTA/AT1G30490, ATHB8/AT4G32880 and CORONA/AT1G52150-as bait to carry out BLAST analyzes (Altschup et al., 1990) using the TBLASTN tool of phytozome (https://phytozome-next.jgi.doe.gov/). Rice and foxtail millet were chosen as monocots representatives, while field mustard, potato and carrot were selected due to their common ability with cassava to form underground storage organs. The rest of the species were selected with the view of encompassing as many vascular plant clades as possible. The only reported C3HDZ sequence of purple anise (*Illicium floridanum*) -belonging to the Amborellales-Nymphaeales-Austrobaileyales (ANA) clade, which contained species are assumed as diverged from the angiosperm common ancestor-was used as outgroup. To generate the phylogenetic tree, we followed the PhyML v3.0 maximum likelihood method (Guindon et al., 2010), making use of the NGPhylogeny.fr server (https://ngphylogeny.fr) (Dereeper et al., 2008). Amino acid sequences in FASTA were used as input (Supplemental File 1). Sequences alignment was performed using the MAFFT v7 tool (Katoh and Standley, 2013), using pre-established parameters. Statistical significance was evaluated through aLRT Likelihood statistics (Anisimova and Gascuel, 2006). Graphic representation of the tree was generated with iTOL v6 (itol.embl.de) (Letunic and Bork, 2021). Synteny analyzes were carried out using the synteny plot tool in Dicots Plaza 5.0 Portal (https://bioinformatics.psb.ugent.be/plaza/versions/plaza_v5_dicots/) (van Bel et al., 2022). Default features and options were used, namely window size: 5, strand orientation and clustering: standard.

### Identification of C3HDZ target promoters expressed in storage roots

We used previously published transcriptomic data to identify genes preferentially expressed in the cassava storage root (Wilson et al., 2017). We then obtained the storage root to fibrous root (SR/FR) expression ratio for all such genes and selected the 245 genes that displayed higher values (Supplemental Table 2). Phytozome v12.1 was used to download, for each gene, up to the first 1000bp of the promoter (i.e: upstream of the first ATG) (Supplemental File 2). We searched for promoters containing canonic C3HDZ binding sites using position frequency matrices (PFM) for *CORONA*, obtained from JASPAR (jaspar.genereg.net) (Castro-Mondragon et al., 2022): a database containing curated, experimentally tested transcription factors binding sites stored as PFMs). Next, we used the webtool MORPHEUS (Minguet et al., 2015) on the generated matrices to analyze the C3HDZ binding site of the 245 sequences. Only promoters containing at least two binding sites in the first 500 bp before the first ATG and obtaining a MORPHEUS score above −10 (which indicates high potential for transcription factor binding) were considered. As an additional validation step for our top candidate promoter, we identified its orthologue Arabidopsis gene and confirmed the presence of C3HDZ binding sites conserved in its promoter using the Promoter Analysais tool from PlantPAN 3.0 http://plantpan.itps.ncku.edu.tw/ (Chow et al., 2018)

### Gene expression and statistical analyses

Cassava plants of 60444 cultivar were grown for 6 months to ensure proper storage root development, as described (Siebers et al., 2017). Storage root, fibrous root, stem, adult leaf and sprout samples were collected. In addition, storage root samples enriched in cambium, xylem or phloem were manually dissected using a razor blade. Given the size of the storage roots, it is very easy to generate such types of samples with a very high degree of accuracy (Siebers et al., 2017). In all cases, samples were immediately frozen in liquid nitrogen and stored at −80°C until needed. RNA was extracted as described (Siebers et al., 2017) and cDNA was generated using the NZY Fist-Strand cDNA Synthesis kit (NZYTech-Genes & Enzymes), following the manufacturers’ instructions. The cDNA was stored at −80°C until used. Primers were designed using the Benchling web tool (www.benchling.com). Used primers are listed in Supplemental table 3. Quantitative RT-PCR reactions were carried out in MicroAmp Fast Optical 96-well reaction plates (Applied Biosystems) using Sybr Green Mix 2x, with ROX as reporter (TakaRa). Three biological replicates were used for each of the above-mentioned sample type. In addition, three technical replicates were performed. Statistical analyses were carried out by using the SPSS 22.0 program (SPSS Inc., Chicago, IL, USA). Mean comparisons were calculated using the Duncan test at 5% probability level.

### Yeast one-hybrid

Bait strains (CZN Manes.06G121400) were generated by transforming YM4271 with the CZN1018 plasmid harboring the *Manes.06G121400* promoter region and selected on SD-Ura medium. The bait strains were transformed with the empty GADT7 plasmid which harbors the GAL4 Activation Domain (AD) (GADT7 AD-Ø) and selected on SD-Ura-Leu to obtain our internal negative controls (CZN Manes.06G121400 + GADT7 AD-Ø). Simultaneously, a subset of the same bait strains was transformed with the GADT7 harboring the MeC3HDZ1 coding region to produce our test strains (CZN *Manes.06G121400* + GADT7 *MeC3HDZ1*). Interaction between MeC3HDZ1 and the *Manes.06G121400* promoter region was examined on SD-Ura-Leu medium supplemented with Aureobasidin A (AbA). The minimal AbA concentration inhibiting growth of the internal negative controls was determined at 150 ng/ml whereas strains showing interaction between the MeC3HDZ1 TF and the promoter region were able to grow on SD-Ura-Leu+AbA medium.

### Luciferase reporter analyses

The first upstream 500bp before the ATG of the promoter region of *Manes.06G121400* were cloned upstream of the *LUCIFERASE* gene in the pGreenII 800-LUC vector (Hellens et al., 2005). In parallel, the *MeC3HDZ1* gene was cloned downstream of 35S:VP16 into the pAlligator1 vector (Bensmihen et al., 2005). The *pManes.06G121400::LUC* construct was co-infiltrated either with the empty pAlligator1 vector *35::VP16* or with the pAlligator1 harboring the *35S::VP16+MeC3HDZ1* construct in *Nicotiana benthamiana* as previously described (Hellens et al., 2005). Luciferase luminescence was quantified in 12 different transfections for each condition (n=12) using a FujiFilm LAS-3000 Luminescent image analyzer as described (Bernabé-Orts et al., 2020).

## Supporting information

Supplemental Fgures

## AUTHOR CONTRIBUTION

JB and JA conceptualized the study and designed the experiments. A.S.-G., A.L, D.O. and C.U. performed the experiments. A.S.-G., A.L, D.O., C.U., J.B. and J.A analyzed, discussed and curated the data. JB and JA wrote the paper. All authors read and approved the final manuscript.

