## Supplemental Fgures for "MeC3HDZ1/MeCNA is a strong candidate for cassava storage root productivity improvement"

### Suplementary Figure 1

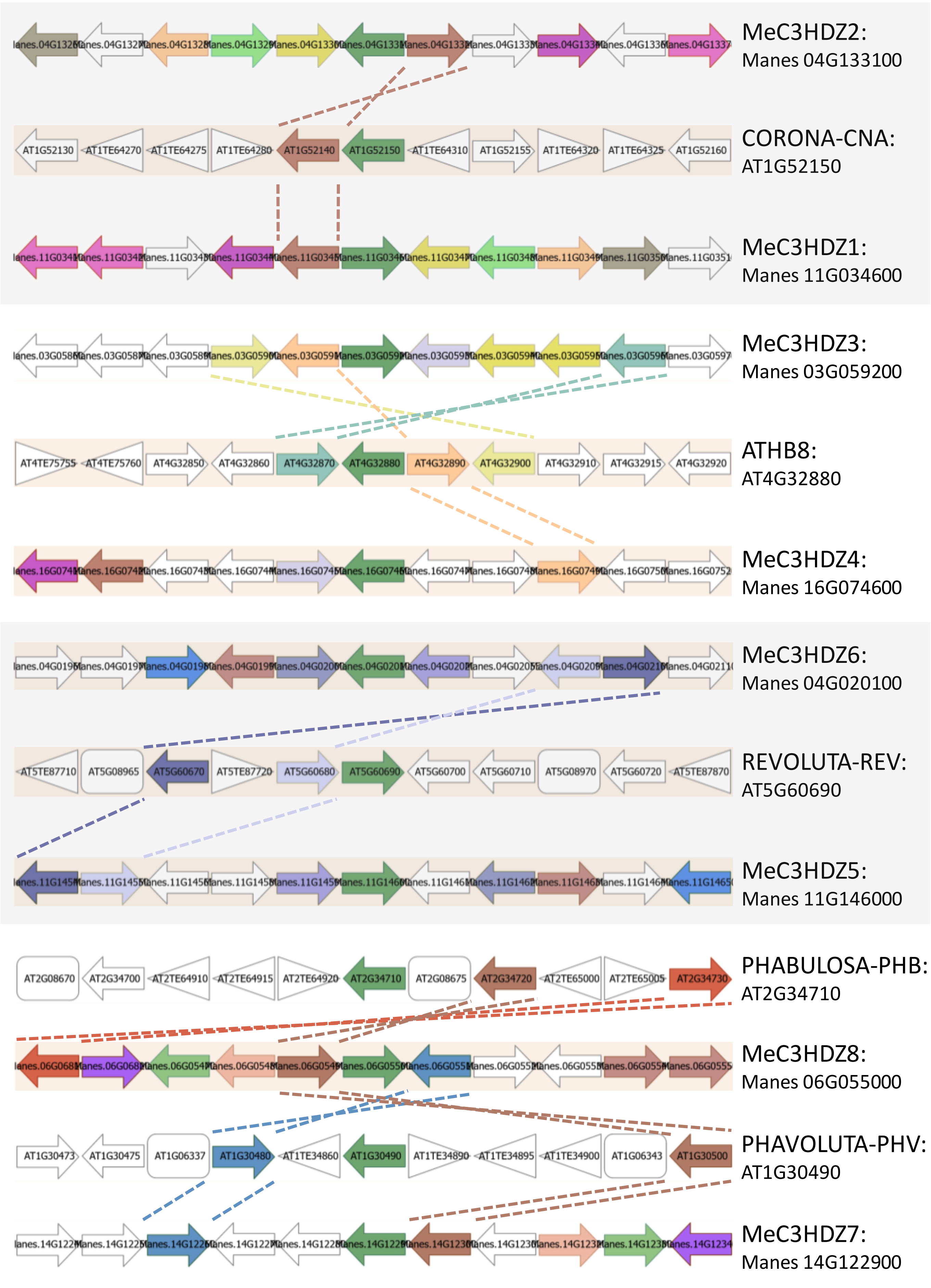

**Supplementary Figure 1.** Synteny analysis of the cassava (*Manihot esculenta*) C3HDZ family employing the synteny plot tool from Dicots Plaza 5.0 Portal. For simplicity, only synteny between cassava and Arabidopsis is shown. Similar results analyses were obtained when analyses were performed performed with 97 species. All examined cassava genes grouped in the *HOM05D000724* gene family

Supplementary Figure 2

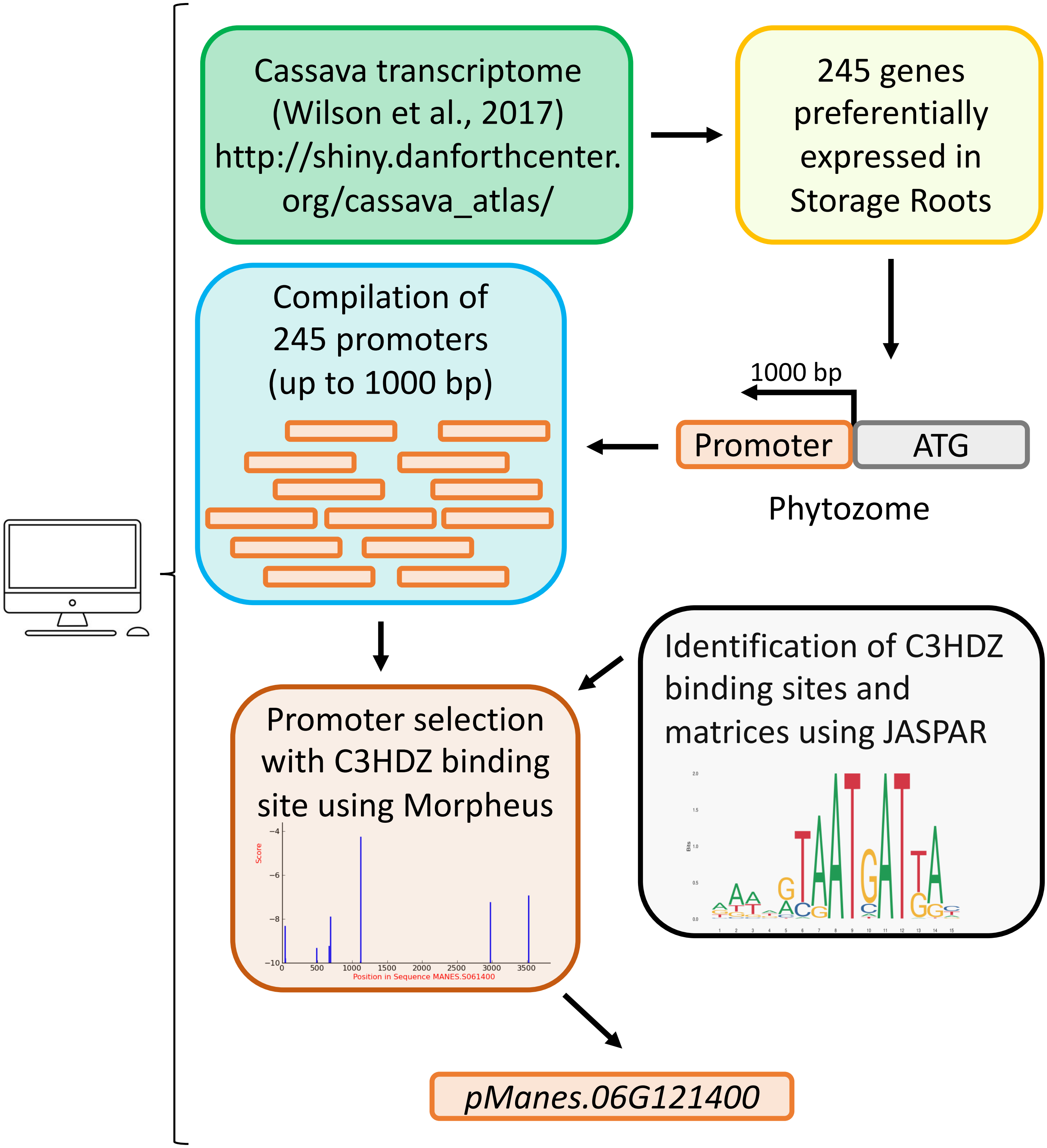


**Supplementary Figure 2.** *In silico* workflow representing how the promoter of *Manes.06G121400* was selected for downstream *in vivo* analysis. We used data from Wilson et al. (2017) to identify genes with strong preferential expression in the storage root (SR). From each of the 245 selected genes, we obtained up to the first 1000bp of the promoter. We used JASPAR (Castro-Mondragon et al., 2012) to identify C3HDZ binding site. Using MORPHEUS (Minguet et al., 2015), we identified promoters with highest probabilities to be bound by C3HDZ, being *pManes.06G121400* our top candidate.

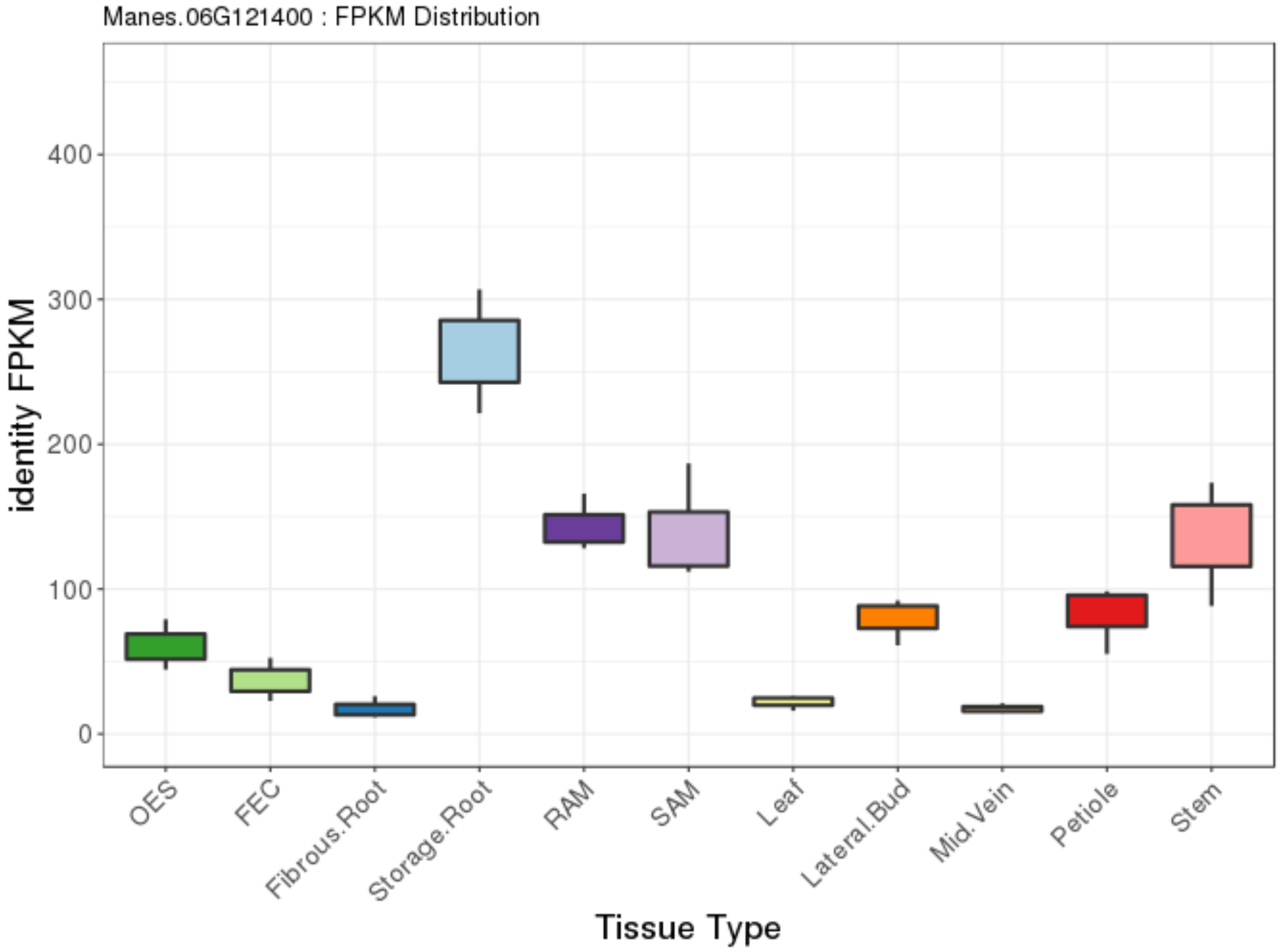

Manes.06G121400 : FPKM Distribution
identity FPKM
400
300
200
100
0
OES
FEC
Fibrous.Root
Storage.Root
RAM
SAM
Leaf
Lateral.Bud
Mid.Vein
Petiole
Stem
Tissue Type


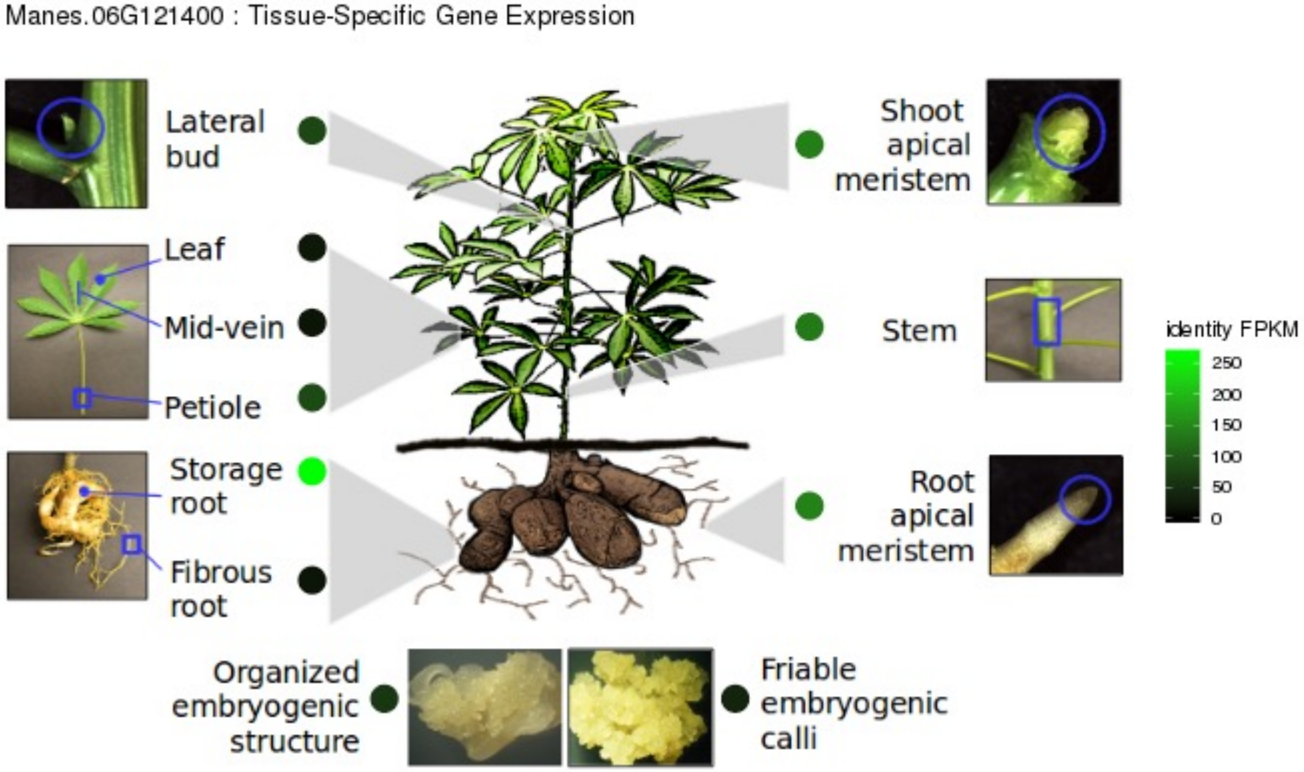

Manes.06G121400 : Tissue-Specific Gene Expression
Lateral bud
Leaf
Mid-vein
Petiole
Storage root
Fibrous root
Organized embryogenic structure
Friable embryogenic calli
Shoot apical meristem
Stem
Root apical meristem
identity FPKM
250
200
150
100
50
0

**Supplementary Figure 3. Expression levels of *Manes.06G121400* are enriched in the storage root*.*** The expression levels of *Manes-06G121400* were checked in previously published transcriptome data of all vegetative organs in cassava (Wilson et al., 2017). Data revealed that the gene is preferentially expressed in the storage root.

A

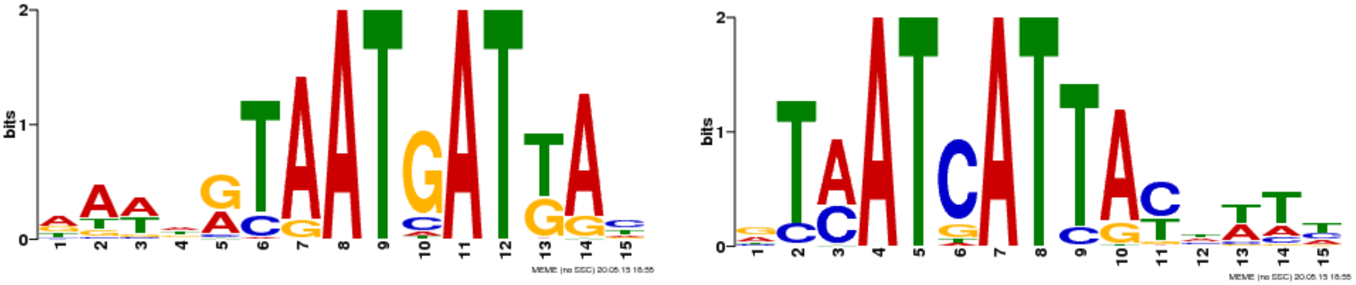

B

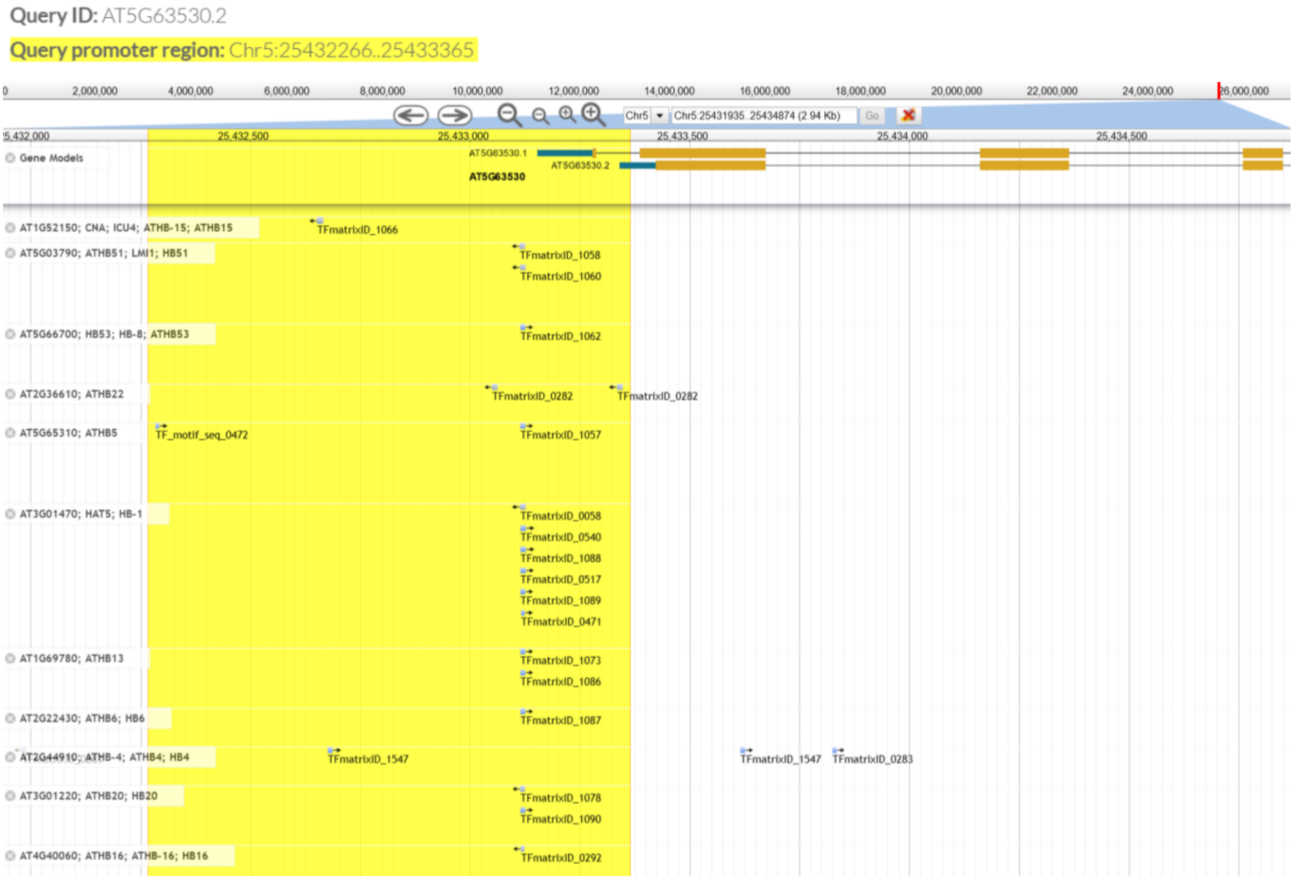


**Supplementary Figure 4. The promoter of *AT5G63530*, the Arabidopsis thaliana orthologue of *Manes.06G121400*, harbors a number of C3HDZ binding sites near its starting transcription site similarly as *Manes.06G121400* promoter**. (A) Memes for the C3HDZ consensus binding sites are shown. (B) The Promoter Analysis tool from PlantPAN 3.0 provides a genome visualization displaying the peaks from ChIP-seq data for several members of the transcription family binding the consensus binding sites on Jbrowse.

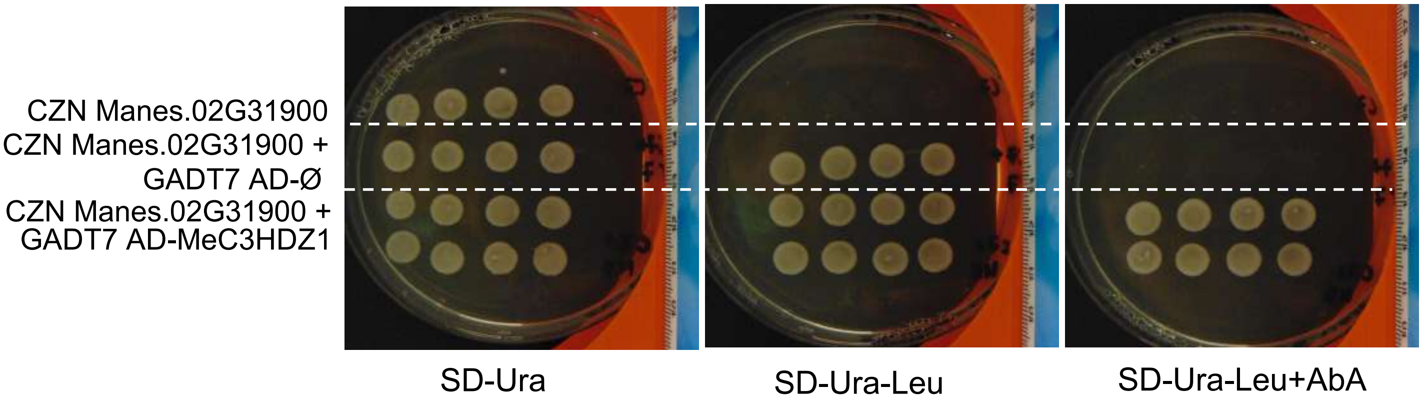


**Supplemental Figure 2. Yeast one-hybrid analyses: original pictures.**
